# Synthetic antibodies neutralize SARS-CoV-2 infection of mammalian cells

**DOI:** 10.1101/2020.06.05.137349

**Authors:** Shane Miersch, Mart Ustav, Zhijie Li, James B. Case, Safder Ganaie, Giulia Matusali, Francesca Colavita, Daniele Lapa, Maria R. Capobianchi, Giuseppe Novelli, Jang B. Gupta, Suresh Jain, Pier Paolo Pandolfi, Michael S. Diamond, Gaya Amarasinghe, James M. Rini, Sachdev S. Sidhu

## Abstract

Coronaviruses (CoV) are a large family of enveloped, RNA viruses that circulate in mammals and birds. Three highly pathogenic strains have caused zoonotic infections in humans that result in severe respiratory syndromes including the Middle East Respiratory Syndrome CoV (MERS), Severe Acute Respiratory Syndrome CoV (SARS), and the ongoing Coronavirus Disease 2019 (COVID-19) pandemic. Here, we describe a panel of synthetic monoclonal antibodies, built on a human IgG framework, that bind to the spike protein of SARS-CoV-2 (the causative agent of COVID-19), compete for ACE2 binding, and potently inhibit SARS-CoV-2. All antibodies that exhibited neutralization potencies at sub-nanomolar concentrations against SARS-CoV-2/USA/WA1 in Vero E6 cells, also bound to the receptor binding domain (RBD), suggesting competition for the host receptor ACE2. These antibodies represent strong immunotherapeutic candidates for treatment of COVID-19.

## INTRODUCTION

Pathogenic strains of coronaviruses have caused serious zoonotic infections in humans three times in the last 20 years. SARS-CoV-2, the cause of the ongoing pandemic of Coronavirus Disease 2019 (COVID-19), is a novel coronavirus belonging to Coronaviridae family, beta-coronavirus genus and Sarbecovirus subgenus. It has a linear, positive-sense, single-stranded RNA genome of approximately 30 kilobases that is 86% identical to that of the SARS-CoV. The SARS-CoV-2 genome shows a high level of homology with SARS-like coronavirus isolated from bats (RaTG13 in particular) and a strain isolated from pangolins (99% identity), and the virus may have originated from recombination between CoVs that infect pangolins and bats (1).

SARS-CoV-2 utilizes an extensively glycosylated spike (S) protein that protrudes from the viral membrane, to mediate host-cell entry (2). The S protein contains 1,282 amino acids organized into 2 subunits (S1 and S2) and forms a homotrimer on the virus surface. The receptor binding domain (RBD), located in the S1 subunit, recognizes the host cell receptor, angiotensin converting enzyme 2 (ACE2), the binding of which facilitates cleavage of the S2 subunit into S2’ and S2” by cell-surface proteases, which in turn enables fusion and internalization of the virus (3). Being responsible for binding to the host ACE2 receptor, the RBD of the S protein presents neutralizing epitopes for COVID-19 (4,5,6) and many studies have shown that the S protein is a major determinant of the immune response in both convalescent patients and in positive asymptomatic individuals.

At the time of writing, there are no approved safe and efficacious therapeutics for treatment or vaccines to protect against COVID-19 available. Several pharmacological treatments are currently under assessment in randomized clinical trials, including many repurposed existing drugs, such as remdesivir, hydroxychloroquine as well as alpha-interferon, and several novel agents targeting key virus-host interactions. Current therapeutic strategies are mainly supportive and aimed at delaying the spread of the virus and to reduce the impact of the disease. Given the severity of the disease, and its rapid spread in the population, more effective therapies and vaccines are needed as soon as possible to deal with this pandemic (7).

To identify antibodies that neutralize SARS-CoV-2 as a basis for therapeutic development, we conducted phage-display selections against the RBD of the S protein, isolated numerous clones that bound to the SARS-CoV-2 RBD or the full-length S protein ectodomain, and screened to identify those that blocked ACE2 binding. In these studies, a library containing 10_10_ Fab-phage clones was used for selections (8). Among the high affinity Fab-phage clones, those that blocked ACE2 were selected for further characterization and expressed as full-length human IgG proteins with a framework engineered to possess high thermostability and low immunogenicity for therapeutic applications (8). Several IgGs were found to exhibit sub-nanomolar affinities for the SARS-CoV-2 S protein and from this group 15033 emerged as a top neutralizer with an ability to neutralize SARS-CoV-2 with high potency.

## RESULTS AND DISCUSSION

We report the identification and characterization of neutralizing synthetic antibodies that bind SARS-CoV-2 RBD. Antibodies identified at the Toronto Recombinant Antibody Centre were isolated by panning phage-displayed Fab libraries against immobilized SARS-CoV-2 S protein receptor binding domain (RBD) in multiple rounds. After 4 rounds of selection, individual clones from selection outputs were amplified as phage supernatants and 384 individual Fab-phage clones were screened by ELISA to compare binding to immobilized RBD and several negative control proteins. Ultimately, 358 clones that bound specifically to the RBD were identified. The CDRs of these antibody clones were decoded by sequencing the DNA encoding the variable regions of each to identify unique clones.

To focus our efforts on clones that bound RBD and blocked its interaction with ACE2, phage ELISAs were conducted on immobilized RBD to compare binding signals on RBD to those on RBD saturated with ACE2. This revealed 38 unique sequences (**Fig. 1A**), whose binding in the presence of ACE2 was inhibited, indicating that they likely bound an overlapping epitope on the RBD. These clones were converted to the full-length IgG format for purification and functional characterization.

**Figure 1.**
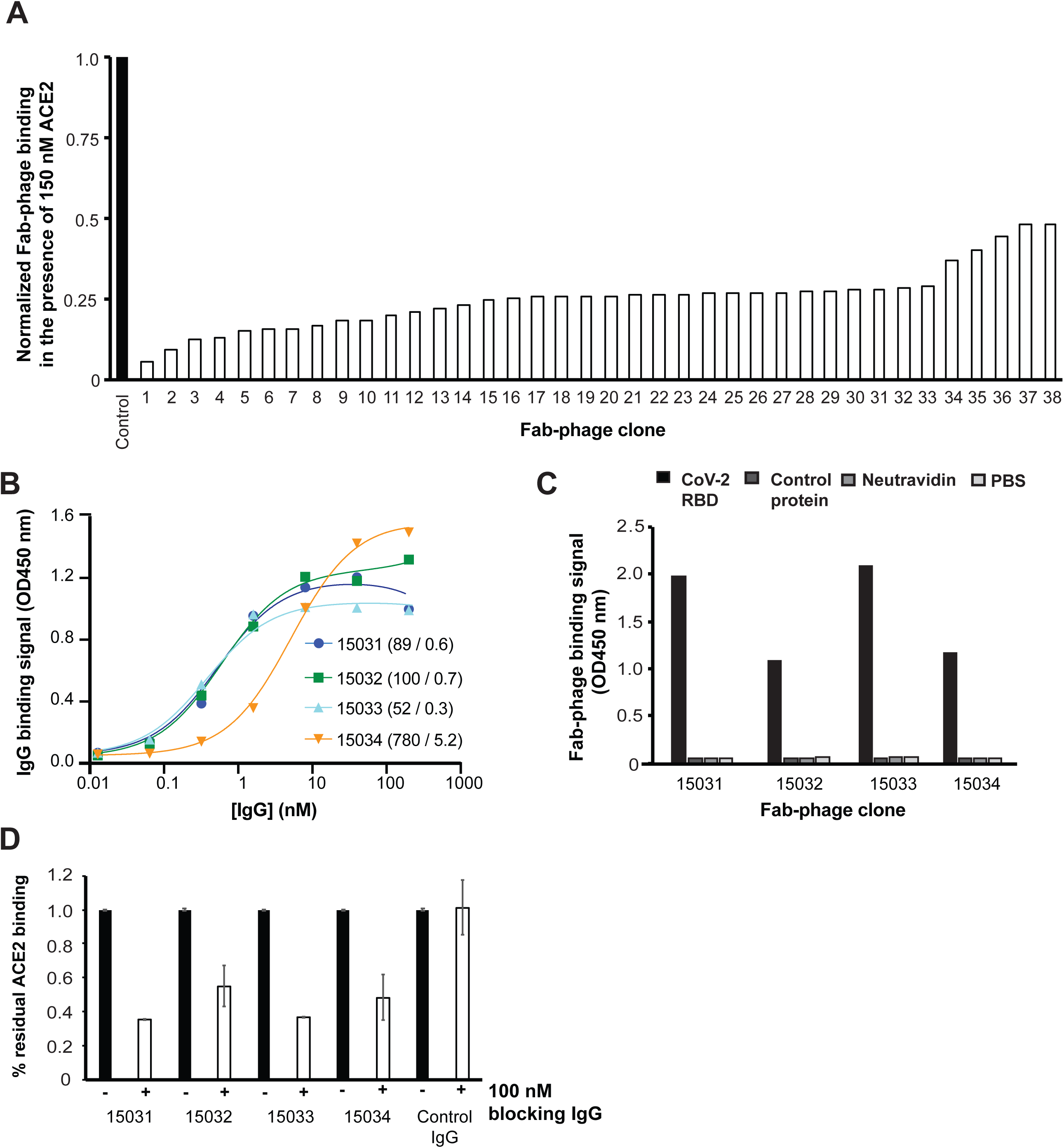
Characterization of IgGs by ELISA. **(A)** Binding of unique Fab-phage clones to immobilized RBD blocked by solution-phase ACE2. Signal was normalized to the signal in the absence of ACE2 (control). (**B)** Serial dilutions of IgGs binding to immobilized SARS-CoV-2 S protein. The EC_50_ values for each curve are shown in parentheses (in ng/mL and nM). (**C**) Binding of Fab-phage to immobilized RBD or negative controls, as indicated. **(D)** Binding of ACE2 to immobilized S protein the presence (white bars) or absence (black bars) of 100 nM IgG.

To estimate affinities, ELISAs were performed with serial dilutions of IgGs binding to biotinylated S protein trimer captured with immobilized neutravidin, and these assays showed that three of four IgGs bound with EC_50_ values in the sub-nanomolar range (**Fig. 1B**). ELISAs also confirmed specificity for RBD, as no signal was observed with immobilized neutravidin alone or with another negative control protein (**Fig. 1C**). To confirm blockade of the interaction between S protein and ACE2 by IgG, the binding of biotinylated ACE2 was assessed by ELISA to immobilized S protein in the presence or absence of saturating IgG; all 4 IgGs inhibited binding (**Fig. 1D**), thus confirming that these antibodies blocked host receptor binding.

To more thoroughly characterize affinities, binding kinetics were assessed by biolayer interferometry (BLI) with biotinylated S protein trimer captured on streptavidin sensors. All 4 IgGs exhibited sub-nanomolar K_D_ values (**Fig. 2**), in close accord with the EC_50_ values measured by ELISA (**Fig. 1B**). In comparing the on and off rates, it is worth noting that IgG 15033 exhibited a k_on_ 2-25-fold faster and IgG 15034 exhibited a k_off_ 5-10-fold slower than those of the other IgGs.

**Figure 2.**
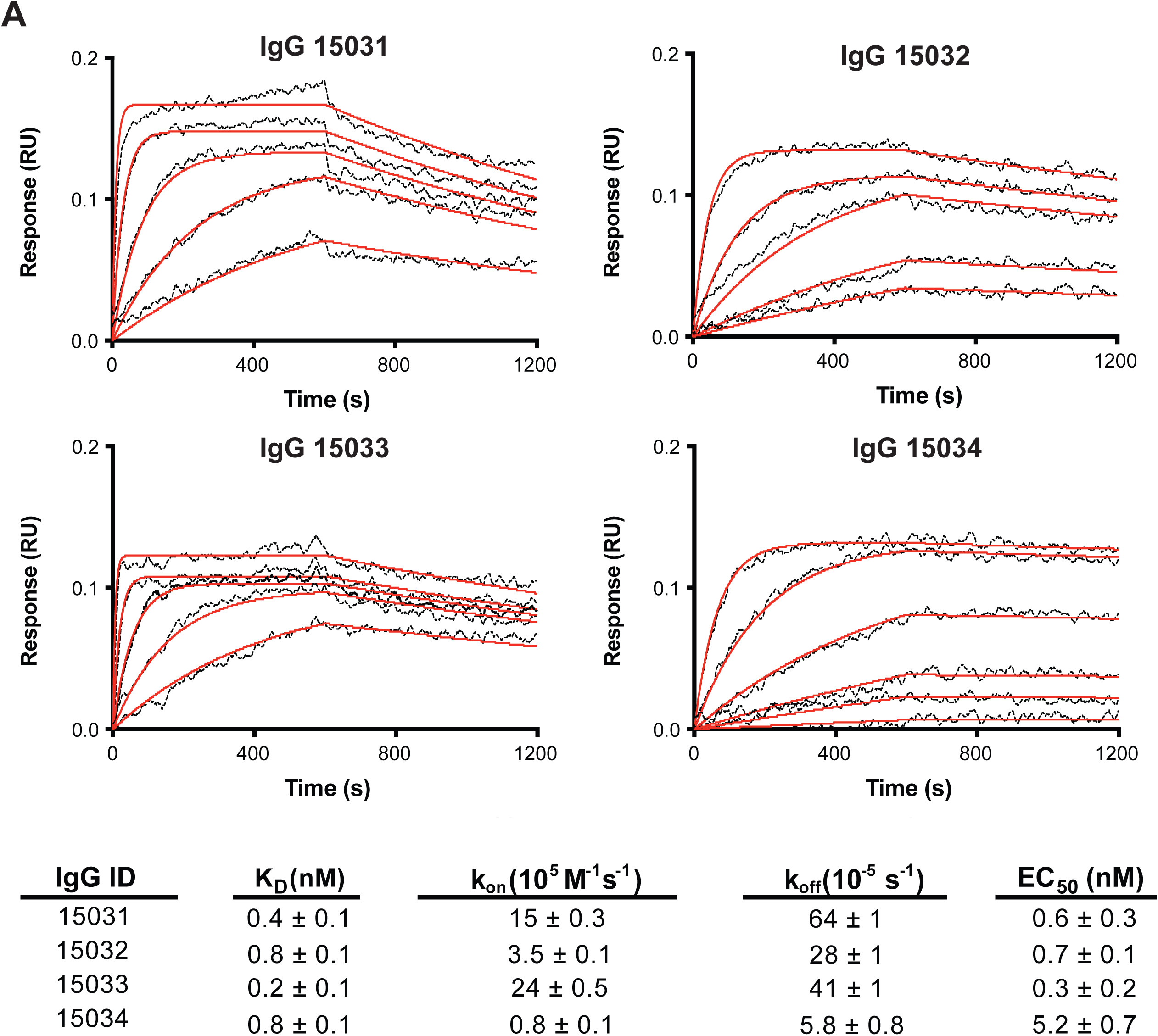
Binding kinetics for IgGs and SARS-CoV-2 S protein. BLI was used to evaluate antibody binding kinetics to immobilized S protein over a range of IgG concentrations (67 - 0.8 nM). Evolved sensor signals (black traces) were fit (red traces) and values for binding kinetics were extracted, and are shown in the table at the bottom, along with EC_50_ values derived from ELISA curves in Figure 1B.

To assess the effects of the IgGs on virus infection, antibodies were evaluated by a microneutralization assay that measured the infection of ACE2-expressing VeroE6 cells with wild-type SARS-CoV-2 (assay performed in St Louis, USA). All 4 IgGs exhibited dose-dependent neutralization of viral infection, confirming their neutralization capacity and the absence of cytopathic effects (**Fig. 3**). All tested antibodies strongly inhibited infection of VeroE6 cells with SARS-CoV-2, with IC_50_ values ranging from ∼100 - 1000 ng /mL (∼0.5-7.5 nM). For the most potent antibody (IgG 15033), inhibition with similar potency (IC_50_ = 80 ng/ml, 0.5 nM) was confirmed in a similar assay at an independent laboratory (assay performed in Rome, Italy). These results confirm that these human antibodies are prime candidates for an anti-viral drug that will block the virus from entering host cells and will thus prevent the virus from replicating and causing disease.

**Figure 3.**
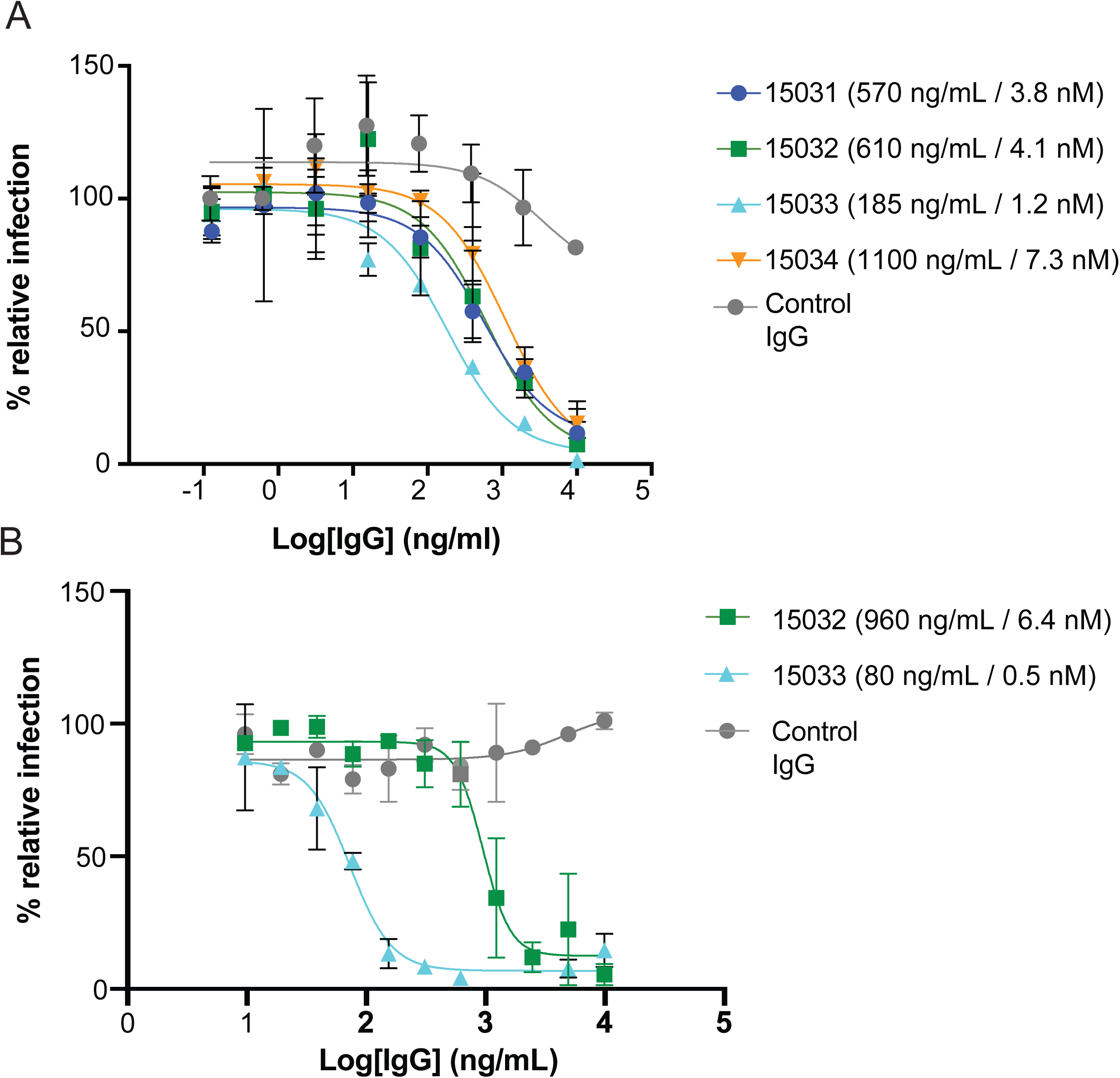
Neutralization of virus by IgGs. Microneutralization assays were performed in St Louis (left) or Rome (right). IgG-mediated neutralization of virus was assessed in a focal reduction neutralization assay and expressed as % relative infection, and infection curves were fitted to extract IC_50_ values, which are shown in parantheses (in ng/mL and nM).

Although therapeutic, antibody-mediated protection against SARS-CoV-2 has yet to be demonstrated in humans, studies suggest that convalescent plasma transfusion may offer benefit to some patients (9–11). Passive virus neutralization appears to not only reduce inflammation, and the lung damage associated with it, but it also works by reducing viral load and preventing dependence on mechanical ventilation. Therefore, therapies with SARS-CoV-2 antibodies must be explored and the mAbs reported here represent valid candidates. Antibodies have the advantages of a long serum half-life and being generally well tolerated, especially if based on a human framework like those described here. In addition to potential therapeutic utility, anti-SARS-CoV-2-specific mAbs are also required for the development of diagnostic and research applications.

## METHODS

### Expression and purification of antigens

The previously reported piggyBac transposase-based expression plasmid PB-T-PAF (12) was modified to generate two new vectors, one containing a CMV promotor (PB-CMV) and the other a TRE promotor (PB-TRE). To facilitate nuclear export of the mRNA, a woodchuck hepatitis virus posttranscriptional regulatory element (WPRE) was also added to each of these plasmids. Each of the protein ORFs was cloned into both expression vectors. The PB-CMV version was used for large-scale transient expression. The PB-TRE version was used for establishing inducible stable cell lines using the piggyBac transposase-based method (12).

cDNAs encoding the SARS-CoV and SARS-CoV-2 spike proteins were human codon optimized and synthesized by Genscript. cDNA encoding human ACE2 was obtained from MGC clone 47598. The soluble SARS-CoV-2 spike ectodomain trimer included residues 1-1211, followed by a foldon trimerization motif (13), a 6xHis tag and an AviTag biotinylation motif (14). Residues 682–685 (RRAR) were mutated to SSAS to remove the furin cleavage site on the SARS-CoV-2 spike protein. Residues 986–987 (KV) were mutated to two proline residues to stabilize the pre-fusion form of the spike (15). The SARS-CoV-2 receptor binding domain (RBD, residues 328-528), the soluble human ACE2 construct (residues 19-615), and the SARS-CoV RBD (residues 315-514) were each followed by a 6xHis tag and an AviTag.

Freestyle 293-F cells were grown in Freestyle 293 expression medium (Thermo Fisher) in suspension culture. Upon transfection, 300 mL cells were seeded to 1-L shaker flasks at a cell density of 10^6^ cells/mL. 15 mL Opti-MEM medium (Thermo Fisher) containing 300 µg of PB-CMV plasmid DNA was mixed with 15 mL Opti-MEM medium containing 400 µL 293fectin reagent (Thermo Fisher). The mixture was incubated for 5 minutes before being added to the shaker flask. Two days post transfection, each 300 mL culture was expanded to 3 shaker flasks containing a total of 900 mL medium, and the expression was continued for another 4 days.

Freestyle 293-F cells or a GnT1-knockout Freestyle 293-F cell line was used for generating the stable cell lines. The cells were transfected on 6-well plates. Each well contained 10^6^ cells growing in 2 mL Freestyle 293 expression medium. For producing doxycycline-inducible cell lines, 2 µg of the PB-TRE expression plasmid, 0.5 µg of the PB-rtTA-neomycin helper plasmid (12) and 0.5 µg of the piggyBac transposase plasmid pCyL43 (16) were co-transfected to each well using the Lipofectamine 2000 reagent (Thermo Fisher). These cells were then selected using 2 µg/mL puromycin and 200 µg/mL G418 for two weeks. The ACE2-expressing cell line was constructed by co-transfecting the Freestyle 293-F cells with the PB-CMV plasmid encoding the full length human ACE2 and the pCyL43 plasmid. This cell line was selected using 2 µg/mL puromycin.

For stable expression, the cell lines were grown as suspension cultures in 1-L shaker flasks. Each flask contained 300 mL Freestyle 293 expression medium supplemented with 1 µg/mL doxycycline and 1 µg/mL aprotinin. Half of the culture was harvested and replaced by fresh medium every other day.

The proteins were purified from the harvested expression medium using Ni-NTA affinity chromatography. The proteins were eluted with phosphate buffered saline containing 300 mM imidazole and 0.1% (v/v) protease inhibitor cocktail (Sigma, P-8849). The proteins were further purified using size-exclusion chromatography. For the RBDs and ACE2, a Superdex 200 Increase (GE healthcare) column was used. For the spike ectodomain, a Superose 6 Increase (GE healthcare) column was used.

Each biotinylation reaction contained 200 µM biotin, 500 µM ATP, 500 µM MgCl_2_, 30 µg/mL BirA, 0.1% (v/v) protease inhibitor cocktail and not more than 100 µM of the protein-AviTag substrate. The reactions were incubated at 30 °C for 2 hours. The biotinylated proteins were then purified by size-exclusion chromatography.

### Fab-phage selections

Fab-phage clones specific for the SARS-CoV-2 S protein were isolated from phage-displayed antibody libraries (8) by multiple rounds of binding selections with the RBD immobilized in wells of neutravidin-coated microwell plates, as described (8). Individual colonies were isolated from *Escherichia coli* infected with phage outputs from rounds 3 and 4, and individual Fab-phage clones were amplified. Antibody variable domains were sequenced by PCR amplification from phage supernatants for extraction of DNA sequences encoding antibody complementarity determining regions.

### Expression and purification of IgG proteins

Phage clone variable domain DNA was amplified by PCR and subcloned into pSCSTa-hIg1 and pSCST1-hk vectors. Vectors for the heavy and light chains were transfected into HEK293F cells (Invitrogen, Grand Island, NY) using FectoPro according to the manufacturer**’**s instructions (Polyplus Transfection, NY). Cell cultures were incubated at 37 **°**C for 4-5 days post-transfection. The cell cultures were centrifuged, and the supernatants were applied to a protein-A affinity column (**∼**2 mL packed beads per 600 mL culture) (Pierce, ThermoScientific, Rockford, IL). IgG proteins were eluted with 100 mM glycine, pH 2.0 and neutralized with 2 M Tris, pH 7.5. The eluent underwent buffer exchange and concentration in to PBS, pH 7.4 by centrifugation in a 50 kDa centrifugal concentrator.

### Binding and competition ELISAs

For capture of biotinylated antigens, 384-well microplate wells were coated overnight at 4 °C with 2 μg/mL neutravidin in PBS pH 7.4. After coating, wells were blocked with 0.2% BSA in PBS for one hour and washed 4 times with 0.05% Tween in PBS (PT buffer). Solutions of S protein (10 nM) or RBD (50 nM) or biotinylated control (50 nM) in PBS were used to immobilize target by incubating for 15 min at room temperature and washing with PT buffer. IgGs were diluted into PBT, applied to the wells, and incubated at room temperature for 30 min. The plates were washed with PBST and incubated for 30 min with anti-k-HRP antibody conjugate (1:7500 dilution in PBT). The wells were washed 4**−**6 times with PBST and developed as described above. The absorbance at 450 nm was determined. To estimate EC_50_ values, data were fit to standard four-parameter logistic equations using Graphpad Prism (GraphPad Software, La Jolla, CA).

For competition ELISAs, S protein was immobilized in 384-well plates as above but at a concentration of 5 **μ**g/well, then wells were quenched with 100 μg/mL biotin. Immobilized S protein was blocked with 100 nM IgG for 30 min and 100 nM biotinylated ACE2 was added to both IgG-blocked and non-blocked wells in parallel. Following a 30 min incubation, the plates were washed with PBST, and HRP/streptavidin conjugate (1:10000 dilution in PBST) was added and incubated for 30 min. The plates were washed with PBST, developed with TMB substrate, and quenched with 0.5 M H_2_SO_4_ before measuring absorbance at 450 nm.

### Binding kinetics

To determine the affinity and binding kinetics of IgGs for the S protein, BLI experiments were performed on an Octet HTX instrument (ForteBio) at 1000 rpm and 25 °C. Biotinylated S protein was first captured on SA biosensors (ForteBio) from a 2 μg/mL solution in PBT, in parallel with an identical concentration of an unrelated biotinylated control protein, followed by a 180 s quench step with 100 μg/mL biotin. After equilibrating with PBT, loaded biosensors were dipped for 600 s into wells containing serial 3-fold dilutions of IgG and subsequently were transferred back into assay buffer for 600 s dissociation. Binding response data were reference subtracted and were fitted with 1:1 binding model using ForteBio’s Data Analysis software 9.0.

### Virus neutralization assays

For assays conducted in St Louis, SARS-CoV-2 strain 2019 n-CoV/USA_WA1/2020 was obtained from the Centers for Disease Control and Prevention (gift of Natalie Thornburg). Virus stocks were produced in Vero CCL81 cells (ATCC) and titrated by focus-forming assay on Vero E6 cells. Serial dilutions of mAbs were incubated with 10^2^ focus-forming units (FFU) of SARS-CoV-2 for 1 h at 37 °C. MAb-virus complexes were added to Vero E6 cell monolayers in 96-well plates and incubated at 37 °C for 1 h. Subsequently, cells were overlaid with 1% (w/v) methylcellulose in MEM supplemented with 2% FBS. Plates were harvested 30 h later by removing overlays and fixed with 4% PFA in PBS for 20 min at room temperature. Plates were washed and sequentially incubated with 1 µg/mL anti-S antibody CR3022 (17) and HRP-conjugated goat anti-human IgG in PBS supplemented with 0.1% saponin and 0.1% BSA. SARS-CoV-2-infected cell foci were visualized using TrueBlue peroxidase substrate (KPL) and quantitated on an ImmunoSpot microanalyzer (Cellular Technologies). Data were processed using Prism software (GraphPad Prism 8.0).

For assays conducted in Rome, SARS-CoV-2 strain 2019-nCoV/Italy-INMI was used. Antibodies were diluted to a concentration of 10 μg/ml in serum-free medium, and titrated in duplicate in two-fold serial dilutions. Equal volumes (50 μl) of approximately 100 TCID50/well virus and antibody dilutions, were mixed and incubated at 37 °C for 30 min. Subsequently, 96-well tissue culture plates with sub-confluent Vero E6 cell monolayers were infected with 100 μl/well of virus-serum mixtures in duplicate and incubated at 37 °C and 5% CO_2_ for two days. Then, the supernatant of each plate was carefully discarded and 120 µl of a Crystal Violet solution containing 2% Formaldehyde was added to each well. After 30 min fixation, the fixing solution was removed and cell viability was measured by photometer at 595 nm (Synergy HTX Biotek).

## REFERENCES

(1) Xiao, K., Zhai, J., Feng, Y., Zhou, N., Zhang, X., Zou, J.-J., Li, N., Guo, Y., Li, X., Shen, X., Zhang, Z., Shu, F., Huang, W., Li, Y., Zhang, Z., Chen, R.-A., Wu, Y.-J., Peng, S.-M., Huang, M., Xie, W.-J., Cai, Q.-H., Hou, F.-H., Chen, W., Xiao, L., and Shen, Y. (2020) Isolation of SARS-CoV-2-related coronavirus from Malayan pangolins. Nature.

(2) Yan, R., Zhang, Y., Li, Y., Xia, L., Guo, Y., and Zhou, Q. (2020) Structural basis for the recognition of SARS-CoV-2 by full-length human ACE2. Science (80-.). 367, 1444–1448.

(3) Hoffmann, M., Kleine-Weber, H., Schroeder, S., Mü, M. A., Drosten, C., and Pö, S. (2020) SARS-CoV-2 Cell Entry Depends on ACE2 and TMPRSS2 and Is Blocked by a Clinically Proven Protease Inhibitor. Cell 181, 271–280.

(4) Wu, Y., Wang, F., Shen, C., Peng, W., Li, D., Zhao, C., Li, Z., Li, S., Bi, Y., Yang, Y., Gong, Y., Xiao, H., Fan, Z., Tan, S., Wu, G., Tan, W., Lu, X., Fan, C., Wang, Q., Liu, Y., Zhang, C., Qi, J., Gao, G. F., Gao, F., and Liu, L. (2020) A noncompeting pair of human neutralizing antibodies block COVID-19 virus binding to its receptor ACE2. Science (80-.). eabc2241.

(5) Shi, R., Shan, C., Duan, X., Chen, Z., Liu, P., Song, J., Song, T., Bi, X., Han, C., Wu, L., Gao, G., Hu, X., Zhang, Y., Tong, Z., Huang, W., Liu, W. J., Wu, G., Zhang, B., Wang, L., Qi, J., Feng, H., Wang, F.-S., Wang, Q., Gao, G. F., Yuan, Z., and Yan, J. (2020) A human neutralizing antibody targets the receptor binding site of SARS-CoV-2. Nature.

(6) Pinto, D., Park, Y.-J., Beltramello, M., Walls, A. C., Tortorici, M. A., Bianchi, S., Jaconi, S., Culap, K., Zatta, F., De Marco, A., Peter, A., Guarino, B., Spreafico, R., Cameroni, E., Case, J. B., Chen, R. E., Havenar-Daughton, C., Snell, G., Telenti, A., Virgin, H. W., Lanzavecchia, A., Diamond, M. S., Fink, K., Veesler, D., and Corti, D. Cross-neutralization of SARS-CoV-2 by a human monoclonal SARS-CoV antibody.

(7) Hung, I. F.-N., Lung, K.-C., Tso, E. Y.-K., Liu, R., Chung, T. W.-H., Chu, M.-Y., Ng, Y.-Y., Lo, J., Chan, J., Tam, A. R., Shum, H.-P., Chan, V., Wu, A. K.-L., Sin, K.-M., Leung, W.-S., Law, W.-L., Lung, D. C., Sin, S., Yeung, P., Yip, C. C.-Y., Zhang, R. R., Fung, A. Y.-F., Yan, E. Y.-W., Leung, K.-H., Ip, J. D., Chu, A. W.-H., Chan, W.-M., Ng, A. C.-K., Lee, R., Fung, K., Yeung, A., Wu, T.-C., Chan, J. W.-M., Yan, W.-W., Chan, W.-M., Chan, J. F.-W., Lie, A. K.-W., Tsang, O. T.-Y., Cheng, V. C.-C., Que, T.-L., Lau, C.-S., Chan, K.-H., To, K. K.-W., and Yuen, K.-Y. (2020) Triple combination of interferon beta-1b, lopinavir–ritonavir, and ribavirin in the treatment of patients admitted to hospital with COVID-19: an open-label, randomised, phase 2 trial. Lancet 395.

(8) Persson, H., Ye, W., Wernimont, A., Adams, J. J., Koide, A., Koide, S., Lam, R., and Sidhu, S. S. (2013) CDR-H3 diversity is not required for antigen recognition by synthetic antibodies. J. Mol. Biol. 425, 803–811.

(9) Li, L., Zhang, W., Hu, Y., Tong, X., Zheng, S., Yang, J., Kong, Y., Ren, L., Wei, Q., Mei, H., Hu, C., Tao, C., Yang, R., Wang, J., Yu, Y., Guo, Y., Wu, X., Xu, Z., Zeng, L., Xiong, N., Chen, L., Wang, J., Man, N., Liu, Y., Xu, H., Deng, E., Zhang, X., Li, C., Wang, C., Su, S., Zhang, L., Wang, J., Wu, Y., and Liu, Z. (2020) Effect of Convalescent Plasma Therapy on Time to Clinical Improvement in Patients With Severe and Life-threatening COVID-19: A Randomized Clinical Trial. JAMA.

(10) Liu, L., Wei, Q., Lin, Q., Fang, J., Wang, H., Kwok, H., Tang, H., Nishiura, K., Peng, J., Tan, Z., Wu, T., Cheung, K. W., Chan, K. H., Alvarez, X., Qin, C., Lackner, A., Perlman, S., Yuen, K. Y., and Chen, Z. (2019) Anti-spike IgG causes severe acute lung injury by skewing macrophage responses during acute SARS-CoV infection. JCI insight 4.

(11) Joyner, M. J., Wright, R. S., Fairweather, D., Senefeld, J. W., Bruno, K. A., Klassen, S. A., Carter, R. E., Klompas, A. M., Wiggins, C. C., Shepherd, J. R. A., Rea, R. F., Whelan, E. R., Clayburn, A. J., Spiegel, M. R., Johnson, P. W., Lesser, E. R., Baker, S. E., Larson, K. F., Ripoll, J. G., Andersen, K. J., Hodge, D. O., Kunze, K. L., Buras, M. R., Vogt, M. N. P., Herasevich, V., Dennis, J. J., Regimbal, R. J., Bauer, P. R., Blair, J. E., Van Buskirk, C. M., Winters, J. L., Stubbs, J. R., Paneth, N. S., and Casadevall, A. Early Safety Indicators of COVID-19 Convalescent Plasma in 5,000 Patients.

(12) Li, Z., Michael, I. P., Zhou, D., Nagy, A., and Rini, J. M. (2013) Simple piggyBac transposon-based mammalian cell expression system for inducible protein production. Proc. Natl. Acad. Sci. U. S. A. 110, 5004–5009.

(13) Tao, Y., Strelkov, S. V., Mesyanzhinov, V. V., and Rossmann, M. G. (1997) Structure of bacteriophage T4 fibritin: A segmented coiled coil and the role of the C-terminal domain. Structure 5, 789–798.

(14) Fairhead, M., and Howarth, M. (2015) Site-specific biotinylation of purified proteins using BirA. Methods Mol. Biol. 1266, 171–184.

(15) Pallesen, J., Wang, N., Corbett, K. S., Wrapp, D., Kirchdoerfer, R. N., Turner, H. L., Cottrell, C. A., Becker, M. M., Wang, L., Shi, W., Kong, W. P., Andres, E. L., Kettenbach, A. N., Denison, M. R., Chappell, J. D., Graham, B. S., Ward, A. B., and McLellan, J. S. (2017) Immunogenicity and structures of a rationally designed prefusion MERS-CoV spike antigen. Proc. Natl. Acad. Sci. U. S. A. 114, E7348–E7357.

(16) Wang, W., Lin, C., Lu, D., Ning, Z., Cox, T., Melvin, D., Wang, X., Bradley, A., and Liu, P. (2008) Chromosomal transposition of PiggyBac in mouse embryonic stem cells. Proc. Natl. Acad. Sci. U. S. A. 105, 9290–9295.

(17) Yuan, M., Wu, N. C., Zhu, X., Lee, C. C. D., So, R. T. Y., Lv, H., Mok, C. K. P., and Wilson, I. A. (2020) A highly conserved cryptic epitope in the receptor binding domains of SARS-CoV-2 and SARS-CoV. Science 368, 630–633.

